# Investigating Medin Cleavage Accessibility in MfgE8: Conformational Insights Derived from Molecular Dynamics Simulations and AlphaFold2 Models

**DOI:** 10.1101/2024.07.27.605412

**Authors:** Shahram Mesdaghi, Rebecca Price, Ming Li, Raymond Q. Migrino, Jillian Madine, Daniel J. Rigden

**Affiliations:** Institute of Systems, Molecular & Integrative Biology, University of Liverpool, Biosciences Building, Crown Street, Liverpool L69 7ZB, UK; Computational Biology Facility, MerseyBio, University of Liverpool, Crown Street, Liverpool L69 7ZB, UK; Phoenix Veterans Affairs, Phoenix, Arizona, USA; University of Arizona College of Medicine-Phoenix, Arizona, USA

## Abstract

Recent studies have indicated that the human amyloidogenic protein medin is associated with a range of vascular diseases, including aortic aneurysms, vascular dementia, and Alzheimer’s disease. Medin accumulates in the vasculature with age, leading to endothelial dysfunction through oxidative and nitrative stress and inducing pro-inflammatory activation. Medin is a cleavage product from the C2 domain of MfgE8. The exact mechanism of medin production from MfgE8 is unknown, with crystal structures of homologous C2 domains suggesting that the cleavage sites are buried, requiring a conformational transition for medin production. Molecular dynamics simulations can explore a wide range of conformations, from small-scale bond rotations to large-scale changes like protein folding or ligand binding. This study employed a combination of full-atom and coarse-grained molecular dynamics simulations, along with CONCOORD- and AlphaFold2-generated models, to investigate MfgE8 conformations and their implications for medin cleavage site accessibility. The simulations revealed that MfgE8 tends to adopt a compact conformation with the RGD motif, important for cell attachment within the N-terminal domain, and the medin region in the C-terminal domain close in proximity. Formation of this compact structure is facilitated by interdomain electrostatic interactions that promote stability and in turn decrease the solvent-accessible surface area of the medin region and particularly the C-terminal medin cleavage site. This data enhances current knowledge on medin generation to propose that alterations in local environmental conditions, possibly through changes in glycosylation or other post-translational modifications are required to induce MfgE8 to unfold partially or fully: this would result in enhanced accessibility of the cleavage sites and therefore enable medin generation.

## Introduction

Increasing evidence shows that medin, a 50-amino acid prevalent but under-studied human amyloidogenic protein, significantly contributes to vascular ageing and age-related cardiovascular and cerebrovascular diseases (Madine et al., 2023). Medin is an internal cleavage product of the milk fat globule-EGF factor 8 protein (MfgE8), comprising positions 268-317 near its C-terminal end (Häggqvist et al., 1999). MfgE8 is a 387-amino acid glycoprotein expressed in various cells - including mammary epithelial cells, endothelial cells, and vascular smooth muscle cells - initially discovered as part of the milk fat globule membrane (Raymond et al., 2009) Reports indicate that medin accumulates in the vasculature with age, leading to endothelial dysfunction through oxidative and nitrative stress, and inducing endothelial pro-inflammatory activation (Davies et al., 2015; Karamanova et al., 2020; Larsson et al., 2006; Migrino et al., 2017; Mucchiano et al., 1992). Medin is linked to the pathophysiology of aortic aneurysms, vascular dementia, and Alzheimer’s disease (Davies et al., 2015, 2019; Migrino et al., 2020). Recent studies have shown vascular medin accumulation in ageing mice, and notably, the selective knockout of the medin-containing domain in MfgE8 preserved cerebrovascular function in aged mice (Degenhardt et al., 2020).

MfgE8 possesses a signal peptide, situated at the N-terminus which plays a crucial role in guiding it to the secretory pathway. MfgE8, also possesses an epidermal growth factor (EGF)-like domain followed by two coagulation domains (C1 and C2) (Figure 1). It is within the N-terminal EGF-like domain that the RGD (arginine-glycine-aspartic acid) motif serves as a critical site for integrin binding (Andersen et al., 1997). The C1 and C2 domains of MfgE8 bind to phosphatidylserine exposed on apoptotic cell membranes, with the C2 domain being the predominant interaction domain (Andersen et al., 1997).

**Figure 1:**
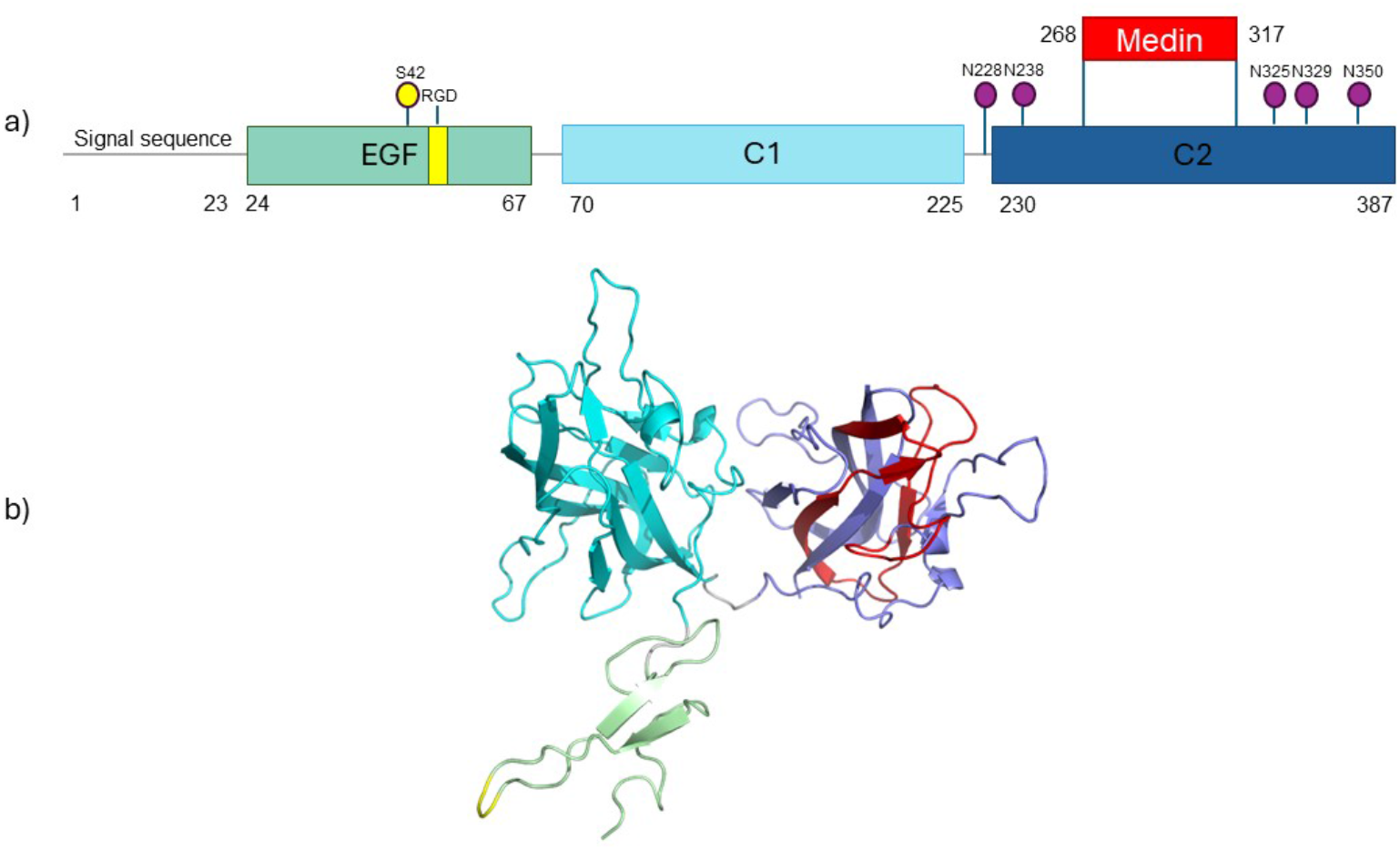
MfgE8 domain and protein structure. a) MfgE8 has a 23 amino acid signal peptide, followed by three distinct domains; EGF, C1 and C2. Identified features of MfgE8 include the RGD motif important for integrin binding (yellow) and medin sequence (red). Post-translational modifications annotated on UniProt (Bateman et al., 2023) include phosphoserine at position S42 (yellow), and glycosylation at asparagine residues at positions 228, 238, 325, 329 and 350 (purple). b) AlphaFold2 model of MfgE8 coloured to correspond to domains, RGD motif and medin region as indicated in part a).

The biological process by which medin is produced from MfgE8 remains unknown, except that it originates through unidentified cleavage mechanisms. Crystal structures of homologous MfgE8 C2 domains indicate that the proteolytic cleavage sites releasing medin are mostly buried and distant from the solvent, suggesting that a conformational transition of the C2 domain is necessary for medin production. Molecular dynamics simulations can uncover these conformational changes, providing valuable insights into this crucial aspect of medin biology. In this study, we utilise a combination of full-atom and coarse-grained molecular dynamics (CGMD) simulations, along with CONCOORD-(CONsensus CONfORMational evaluation of COORDinates) (de Groot et al., 1997) and AlphaFold2 (Jumper et al., 2021) -generated models, to explore the conformations that MfgE8 adopts and how these conformations impact the accessibility of the putative medin cleavage sites.

## Methods

The MfgE8 protein structure model was obtained from the AlphaFold2 Protein Structure database (Varadi et al., 2022). PyMOL (Delano, 2002) was used to truncate the MfgE8 structure, removing the first 23 residues corresponding to the signal peptide which was predicted by submitting the sequence to the SignalP server (Almagro Armenteros et al., 2019).

Structures were solvated with the TIP3P water model (Jorgensen et al., 1983). The system was neutralised with Na^+^/Cl^-^ ions in order to generate the conditions for computing the long-range electrostatic interactions, which requires charge neutrality of the system. Long-range electrostatics were calculated using particle mesh Ewald algorithms (Pronk et al., 2013). Energy minimization was conducted for up to 10,000 steps, utilising first the steepest descent algorithm and then the conjugate gradient algorithm. The system was then heated from 0 K to 300 K. After two equilibration steps, each lasting 20 ps, simulations were run for 1 µs with a 2 fs timestep. The three replicate 1μs MD simulations were performed in GROMACS 2022.2 (Van Der Spoel et al., 2005) molecular dynamics simulation software and carried out using the CHARMM36 (Lindahl et al., 2010) force field. Three replicate MD simulations were carried out to improve the reliability of results. Visualisation of the results was performed with VMD 1.9.3 (Humphrey et al., 1996). All molecular dynamics analysis was conducted using Python scripts with the MDAnalysis (Michaud-Agrawal et al., 2011) and MD-DaVis (Maity & Pal, 2022) modules.

In order to determine whether the compact structure is a preferred conformation a large number of coarse-grained simulations were run; ten replicates starting with the AlphaFold database model and ten replicates using the compact model obtained from the last frame of the first all atom trajectory. First, using the publicly available python script (martinize.py) and a local installation of DSSP (Bulut & Korpeoglu, 2007), the models were converted from an atomistic protein structure into a coarse-grained mode where four heavy atoms are grouped together in one coarse-grain bead. Each residue has one backbone bead and zero to four side-chain beads depending on the residue type. Twenty 1μs coarse-grain replicate simulations were performed using GROMACS 2022.2 molecular dynamics simulation software with Martini v2.2 (Siewert J. Marrinket al., 2007) force field.

All simulations were performed on an Ubuntu 20.04.6 workstation AMD Ryzen Threadripper 2990WX 32 Core CPU (3.0 GHz) with 64GB RAM. GPU acceleration was performed by an ASUS TUF GeForce RTX 3080 OC LHR 12GB GDDR6X Ray-Tracing Graphics Card, 8960 Core, 1815MHz Boost.

PISA (protein interfaces, surfaces and assemblies) (Krissinel & Henrick, 2007) was used to analyse the interfaces of the interacting domains to determine whether there are specific residue contacts responsible for the interaction of the different domains.

Consurf (Ashkenazy et al., 2016) was used to map residue conservation scores onto the MfgE8 model and co-variance analysis using ResPre (Li et al., 2019) was used to predict contacts at the interdomain interacting interface. For both tools initially a multiple sequence alignment (MSA) was generated using HMMER (Potter et al., 2018) to identify similar sequences in UniProt (Bateman et al., 2023)which were then filtered to remove those that were too divergent or redundant, ensuring coverage of the region of interest. Due to contamination of the MSA with alignments against other proteins containing EGF-like and C1/2 domains, HMMER was secondarily used to identify similar sequences, and an MSA containing the closest 100 sequences to the input was used for Consurf mapping of residue conservation.

The solvent-accessible surface area (SASA) was computed using VMD (Humphrey et al., 1996). SASA was assessed for the entire medin region, as well as using 3-residue, 10-residue, and 1-residue windows. GROMACS software was employed to determine the root mean square deviation (RMSD) and radius of gyration (Rg).

The C1 and C2 sequence alignments were performed by HHAlign as implemented in the HHPred server (Gabler et al., 2020) with default parameters. The structural alignment of these domains (obtained from the AlphaFold database) was carried out with the DALI server (Holm, 2022).

## Results

### Trajectory 1 shows that MfgE8 adopts a compact conformation

The initial AlphaFold database (AFDB) model (Figure 1b) was subjected to three replicate 1μs MD simulations. The root mean square deviation (RMSD) of each frame with respect to the reference structure (the initial AFDB model) was measured. Inspection of this RMSD profile for the first replicate shows two distinct phases; in the first 0.3μs the RMSD fluctuates at around 2.55Å, followed by a jump in RMSD to around 2.7Å with more consistency in the readings thereafter (Figure 2).

**Figure 2:**
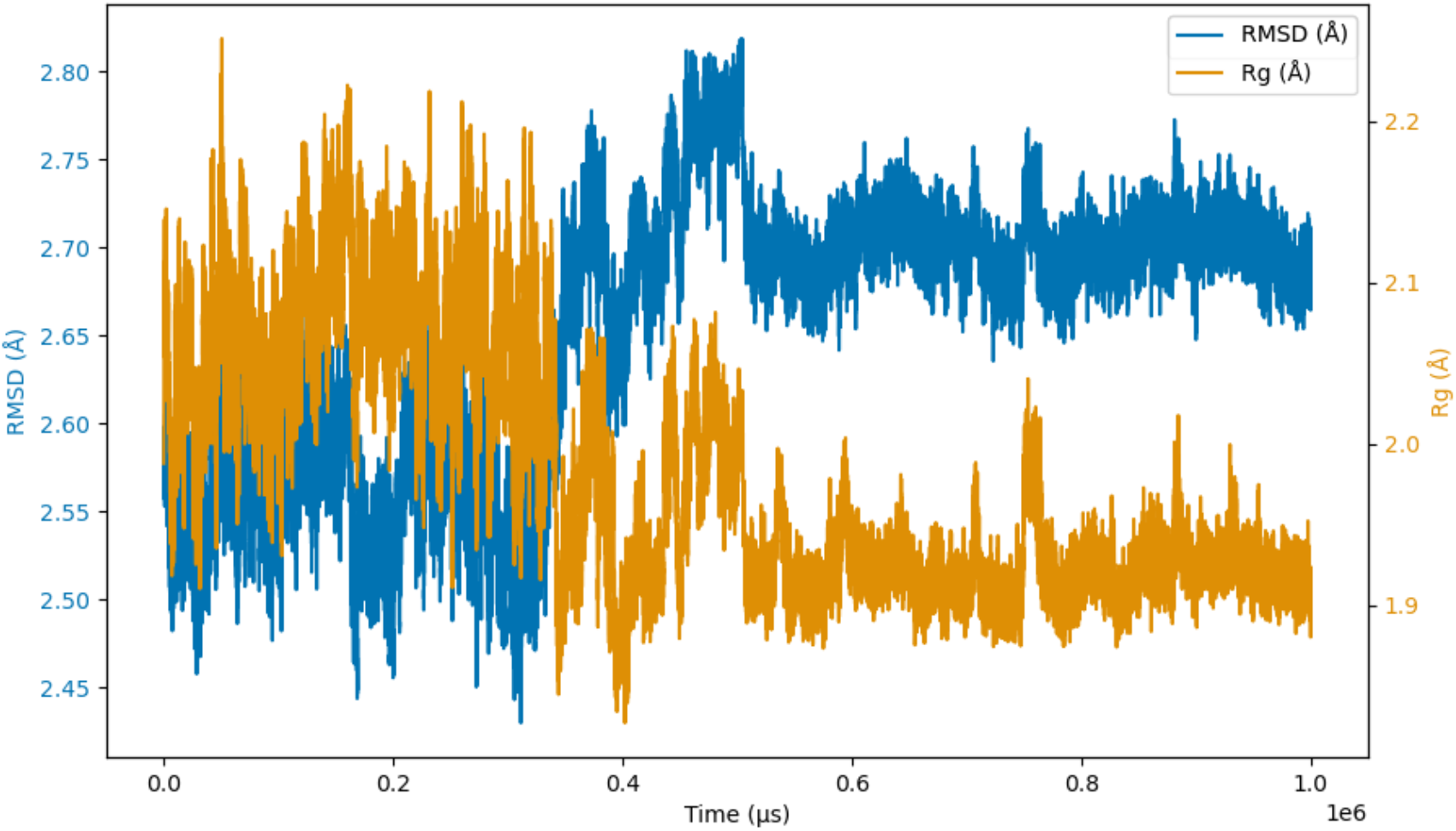
Replicate 1 RMSD and Rg profile. The profile reveals two distinct phases: during the initial 0.3 μs, the RMSD fluctuates around 2.55 Å, followed by an increase to approximately 2.7 Å, with more consistent readings observed thereafter. Similarly, there is a drop in Rg at the 0.3μs mark with more stable readings in the Rg thereafter.

The radius of gyration (Rg) is a measure of compactness of a structure. The values were used to build a profile illustrating how the compactness of MfgE8 changed over time during the simulations. Examination of the Rg profile for the first replicate clearly shows a drop in Rg at the 0.3μs mark with more stable readings in the Rg thereafter (Figure 2). Cross-referencing the Rg profile with the RMSD profile shows that their shifts coincide around the same point in the trajectory.

The free energy of a protein is directly related to protein stability. GROMACS was used to calculate the free energy value for MfgE8 in each frame of the first replicate. These values were then used to plot a free energy landscape in terms of RMSD and Rg. The protein free energy plot visualises the protein’s conformational states and their associated energies. Minima on the plot represents stable conformational states, with the global minimum corresponding to the most stable. Inspection of the landscape indicates that the shift in RMSD and Rg in the latter half of the simulation corresponds to a free energy minimum (Figure 3). The minima in free energy on a free energy landscape are assumed to correspond to stable conformations of the protein.

**Figure 3:**
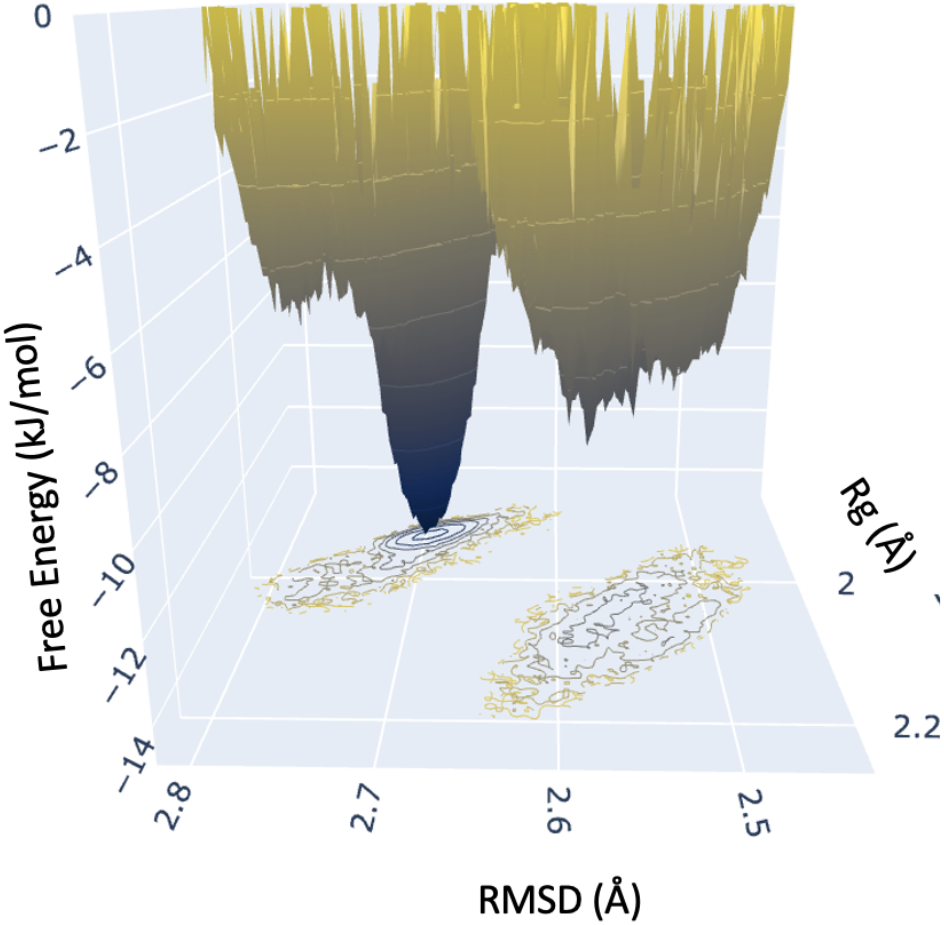
Replicate 1 Free energy landscape. The energy minimum corresponds to a global minimum in free energy, suggesting that the protein is more stable and less likely to undergo spontaneous conformational changes in this state - the stability arises because the protein has reached a point where the potential energy is lower compared to other conformations so the protein is more likely to reside in this conformation for longer periods. The landscape analysis reveals that the shift in RMSD and Rg observed in the latter half of the simulation corresponds to a free energy minimum.

### The medin region interacts with the EGF-like domain RGD motif

Examination of the surface electrostatics of MfgE8 revealed that there is a negative region on the EGF domain and a positive patch on the C2 domain (Figure 4a). Visualisation of the coordinate files from the first replicate trajectory show that the energy minimum corresponds with an electrostatic interaction of the C-terminal side of the C2 medin-containing domain with the EGF-like domain. Interestingly, this region contains the RGD (46-48) motif responsible for cell adhesion and integrin binding (Figure 4b,c).

**Figure 4:**
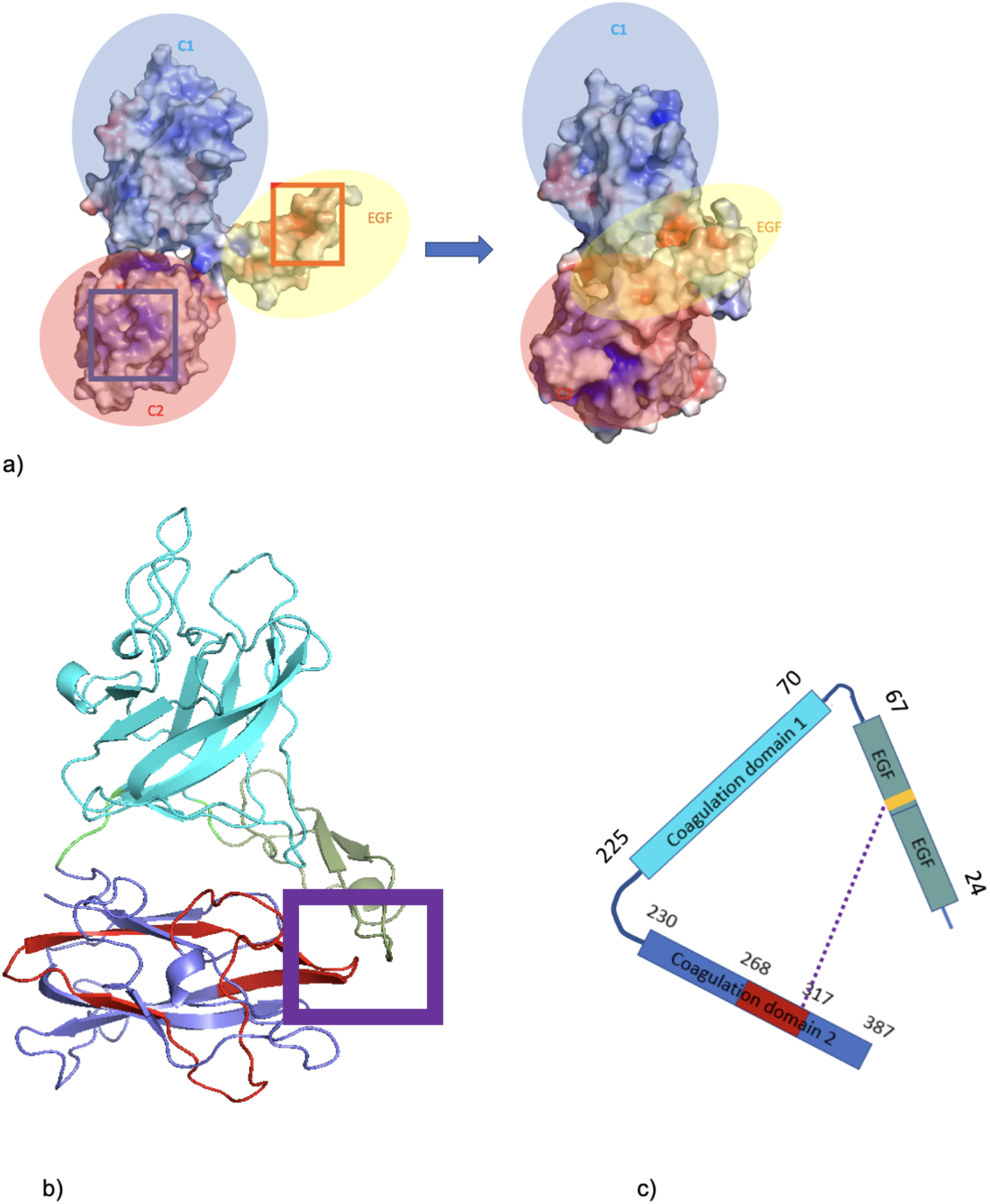
Evaluation of compact structure formed in replicate 1 trajectory. a) MfgE8 transitioning from the AFDB conformation to the compact state with surface electrostatics mapped. Red (negative) to Blue (positive). Red box indicates anionic patch on the EGF-like domain that interacts with the C2 cationic region (blue box). b) Structure of MfgE8 taken from the final frame of replicate 1. Red indicates medin region. Purple box highlights the interaction of the C2 domain with the EGF-like domain. c) Schematic of MfgE8. Purple dotted line indicating the RGD motif interaction with the C-terminal side of the C2 medin containing domain. Cyan is the C2 domain; blue is the coagulation domain 2; green is the EGF-like domain; red is the medin region; yellow is the RGD motif.

Furthermore, hydrogen bond analysis reveals the presence of hydrogen bonds between the C2 medin-containing domain and the EGF-like domain (Figure 5) in the latter part of the simulation.

**Figure 5:**
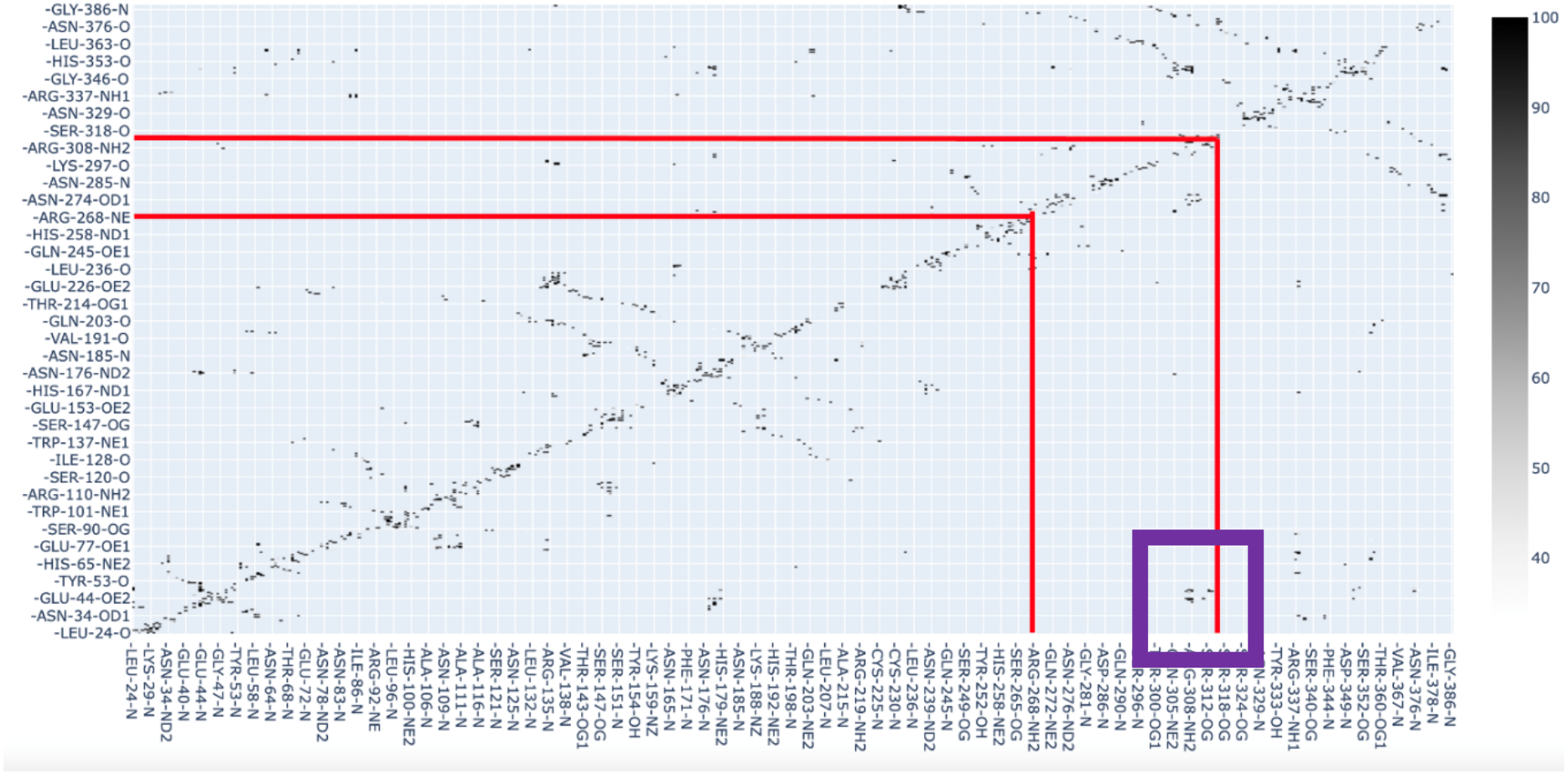
Hydrogen bond matrix for replicate 1. Each point is a hydrogen bond contact, the darker the point the more frames the contact was present (see spectrum on right). Red lines indicate medin boundaries. Purple box highlights the contacts between C2 medin-containing domain with the EGF-like domain.

The distance profile of the β-carbons (the first carbon atom of the sidechain in an amino acid) from R308 (within the medin region of the C2 domain) and D48 (of the RGD motif) over the 1μs replicate 1 simulation shows how the dramatic conformational shift brings the RGD motif and the medin region (C-terminal side) together (Figure 6) in a in a three-dimensional structure which is stabilised by hydrogen bond interactions.

**Figure 6:**
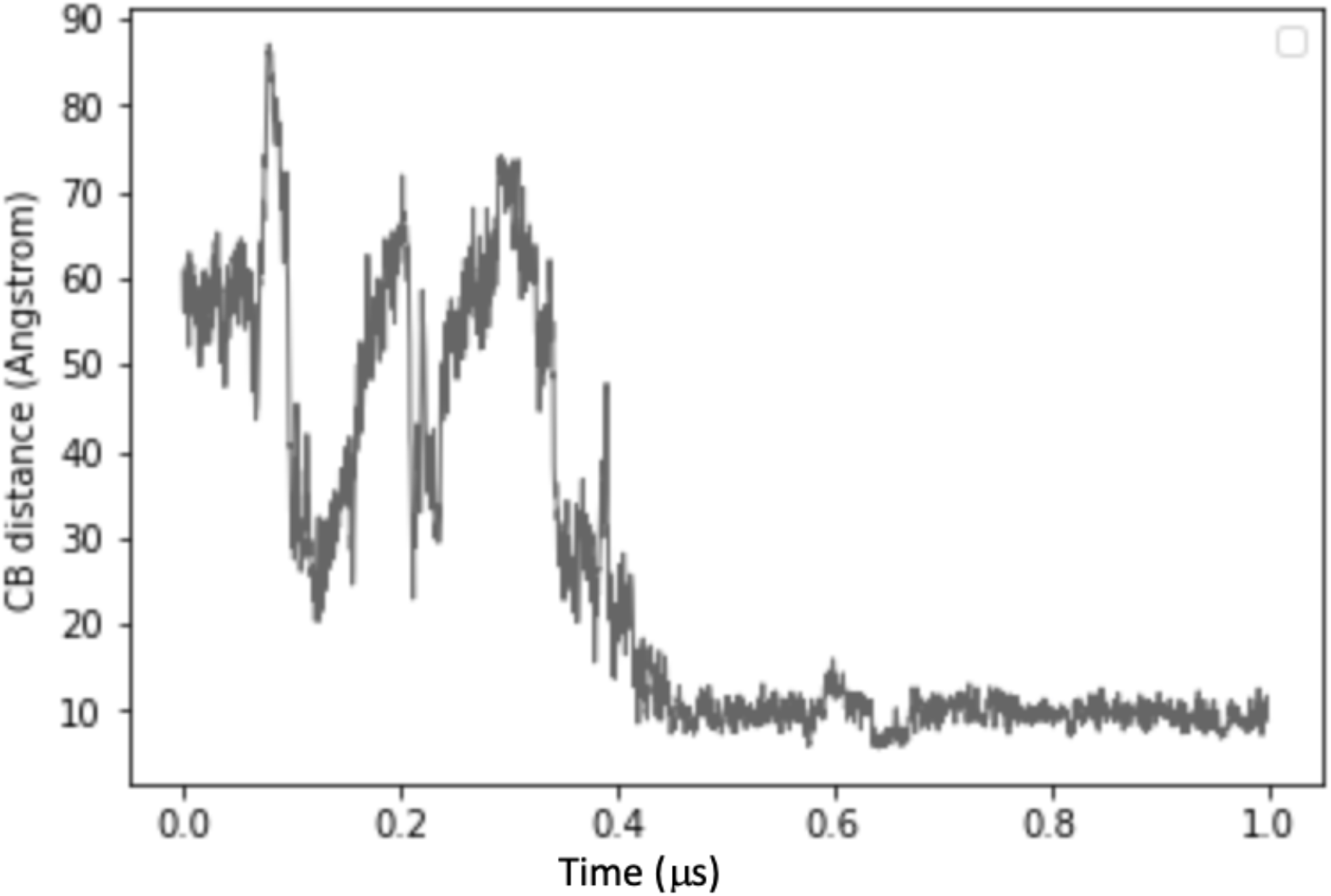
R308 and D48 Cβ distances over replicate 1 trajectory.

Further analysis of residue interactions at the inter-domain interfaces using PISA across the entire trajectory for the first replicate did not reveal any additional interactions (data not shown). Covariance analysis did not identify any predicted contacts corresponding to observed structural contacts, thereby not supporting the presence of specific contacts that are responsible for maintaining the compact conformation (data not shown).

### Coarse-grained simulations reveal MfgE8 can adopt different compact conformations

The second and third replicate 1μs MD simulations did not show the dramatic shift into a stable compact conformation that arrives at a free energy minimum observed in the first replicate (Figure 7).

**Figure 7:**
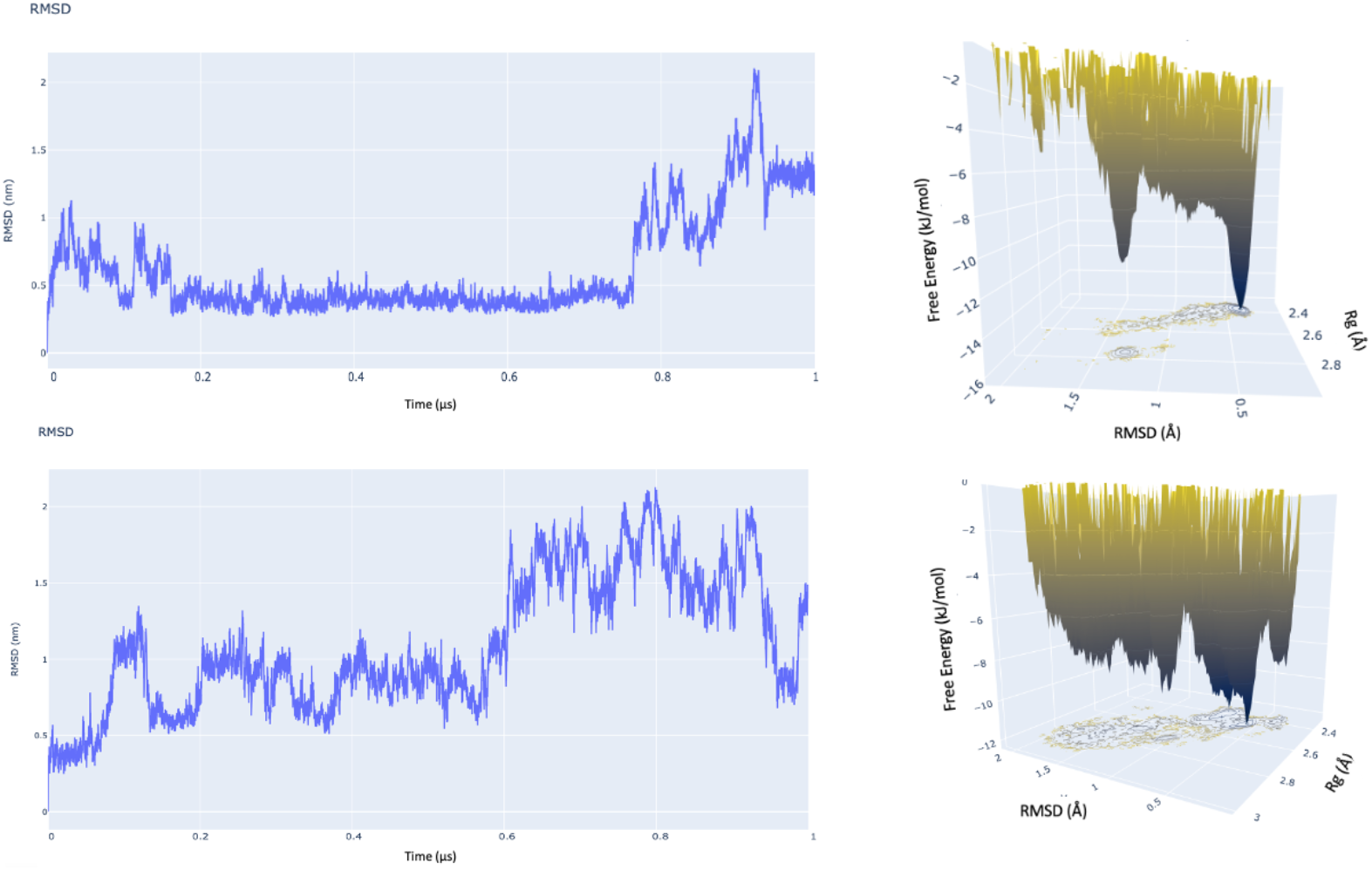
RMSD and free energy plots for replicates 2 (top) and 3 (bottom). In contrast to replicate 1, no stable conformation was achieved in these replicates.

Due to local fluctuations and collective motions occurring simultaneously it is hard to distinguish the two types of motion from each other: therefore, principal component analysis (PCA) was used to filter global, collective (often slow) motions from local, fast motions. A set of eigenvectors and eigenvalues were calculated, using the three trajectories concatenated, which describe collective modes of fluctuations of MfgE8. The eigenvectors corresponding to the largest eigenvalues represent the largest-amplitude collective motions and the top three eigenvectors were used as conformational descriptions. The analysis indicated that the three trajectories shared little conformational space (Figure 8).

**Figure 8:**
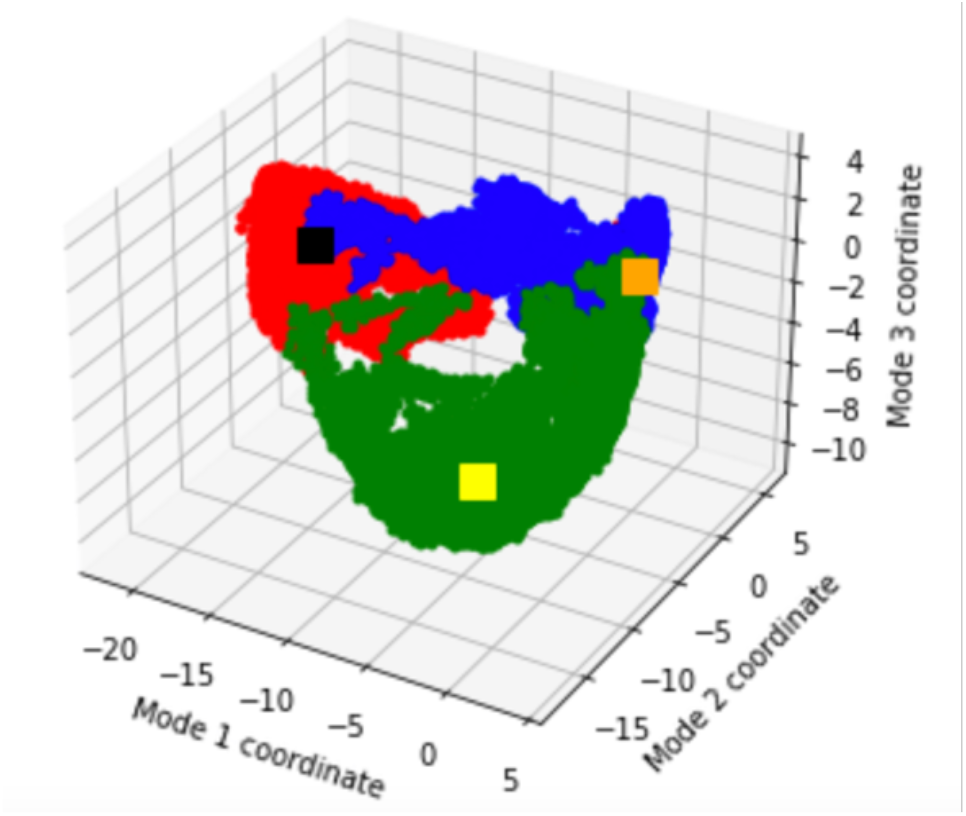
Conformational space occupied by the three replicates in terms of their top three modes. Red - trajectory 1, black square is the final point of trajectory 1. Blue – trajectory 2, yellow square is the final point of trajectory 2. Green – trajectory 3, orange square is the final point of trajectory 3.

In an effort to evaluate whether the compact conformation is the preferred state of MfgE8, a large number of coarse-grained simulations were run; ten replicates starting with the AFDB model and ten replicates using the compact model obtained from the last frame of the first replicate all atom trajectory. Using GROMACS the distances between the centroids (centre of geometry) of the EGF domain and the C2 domain was calculated for the duration of the simulations. In 19 out of the 20 trajectories MfgE8 transitions into a compact conformation (Figure 9). The distance traces indicate that even though they fall into a compact conformation, they do not fall into the same compact conformation.

**Figure 9:**
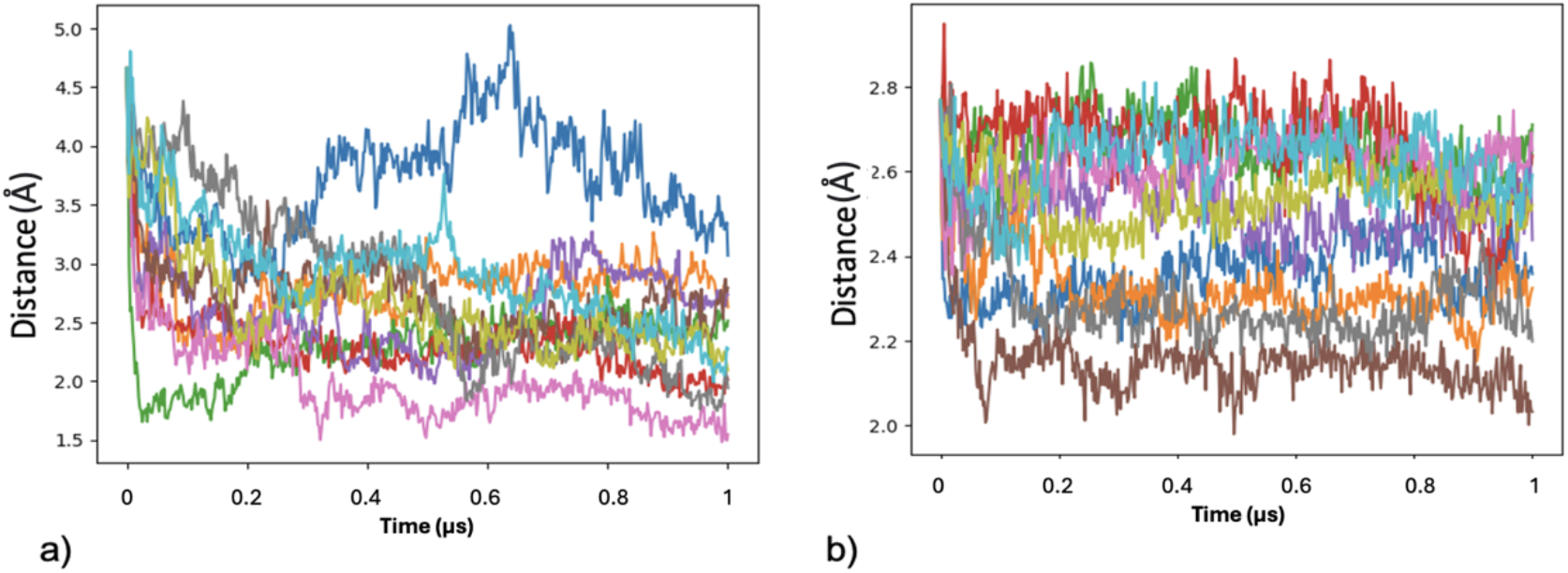
C2 and EGF domain distances over 1μs. a) 10 trajectories starting with the MfgE8 AFDB model. b) 10 trajectories starting with the compact MfgE8 model from replicate 1.

ConSurf mapping was employed to identify conserved residues at identified interface regions in compact structures generated. Mapping the residue conservation scores onto the MfgE8 model indicated a conserved region spanning residues 300-311, incorporating the R308 residue identified in the replicate 1 compact conformation described in detail above (Figure 10a). Electrostatics surface mapping of the AFDB C2 domain model identified this region to be positively charged, consistent with electrostatic interactions driving the potential for compact conformation formation with the RGD motif region (Figure 10b).

**Figure 10.**
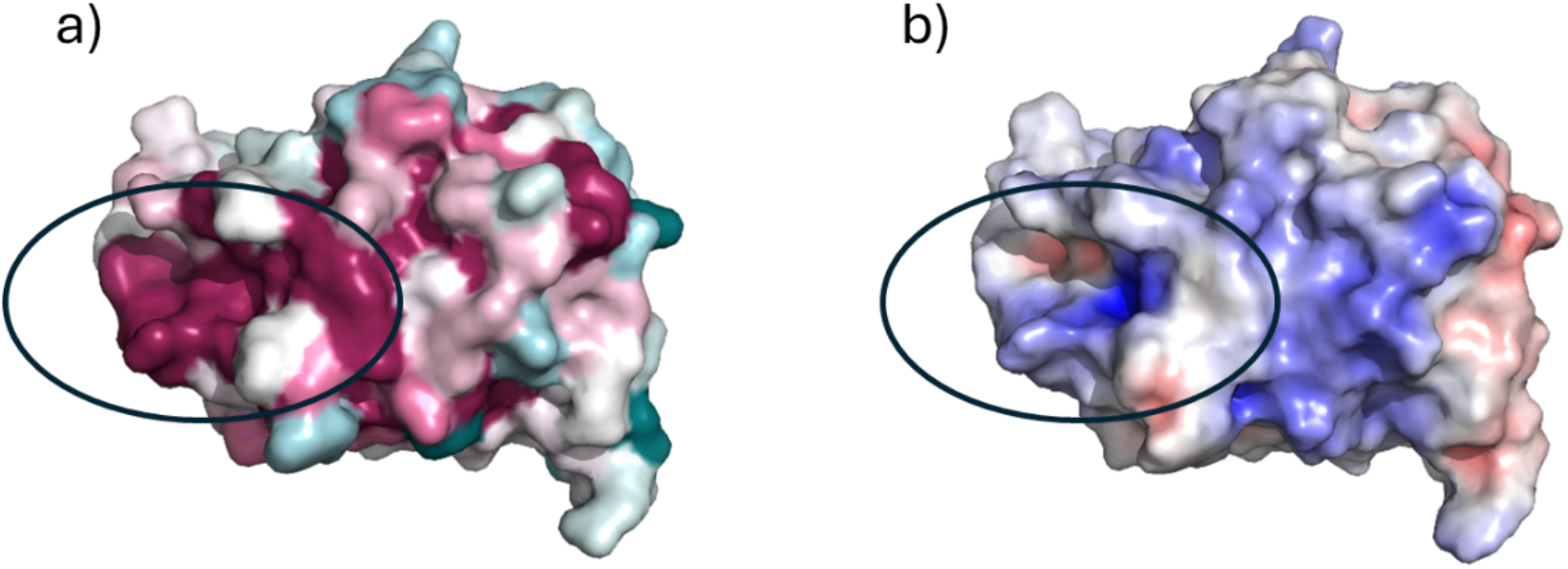
AFDB model of the MfgE8 C2 domain. a) Consurf surface view (pink – conserved to green - low conservation) revealing a highly conserved region (residues 300-311) and b) Electrostatics surface mapping (negative – red to positive – blue) revealing a positively charged region corresponding to the conserved residues.

### MfgE8 cleavage site accessibility is minimally impacted by conformational shifts

To assess the accessibility of the medin cleavage site, we conducted Solvent Accessibility Surface Area (SASA) analyses on structures derived from both all-atom and coarse-grained simulations. Analysis of the average SASA per residue indicates that the cleavage sites reside within regions of relatively low SASA (see Figure 11). This observation holds consistently across all three replicates of the all-atom simulations and the entire set of coarse-grained simulations, despite divergence in their explored conformational spaces. For the AFDB model, SASA values at the potential medin cleavage sites were measured at 550Å^2^ for the N-terminal and 250Å^2^ for the C-terminal regions. In replicate one, SASA values ranged from 400 to 600 at the N-terminal cleavage site and from 150Å^2^ to 300Å^2^ at the C-terminal site.

**Figure 11:**
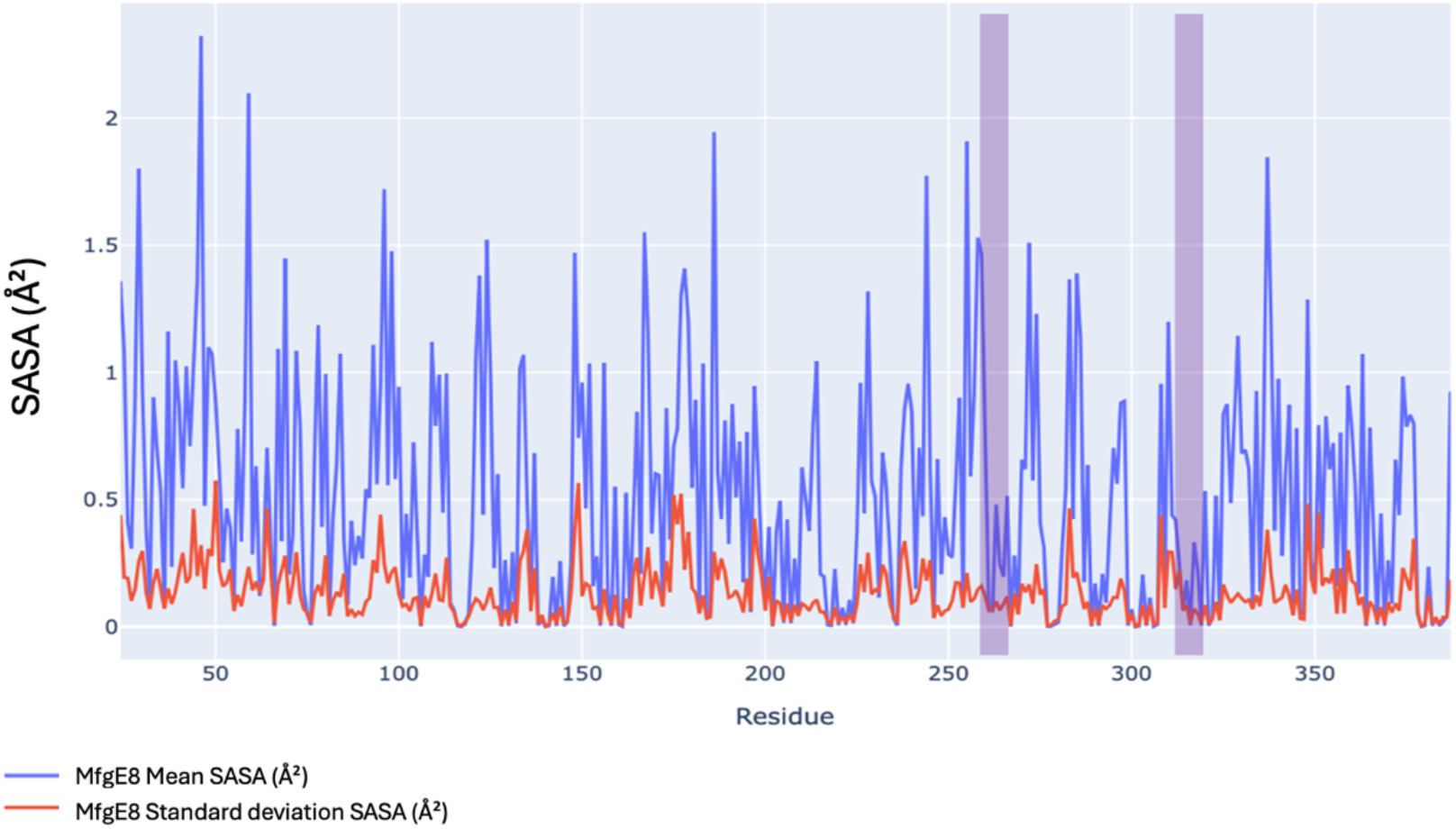
Mean and standard deviation of SASA per residue profile for all atom replicate 1. Blue trace is the mean SASA and the red trace is the SASA standard deviation. Purple highlights the position of the putative medin cleavage sites.

Investigation of SASA profiles throughout the 1 μs trajectories shows that the SASA across the medin region decreases when MfgE8 adopts a compact conformation (Figure 12a and f). Further analysis of 10- and 3-residue windows surrounding the cleavage sites indicate that the same trend of decreasing accessibility upon formation of the compact conformation is only observed for the C-terminal 10-residue analysis (Figure 12b-e).

**Figure 12:**
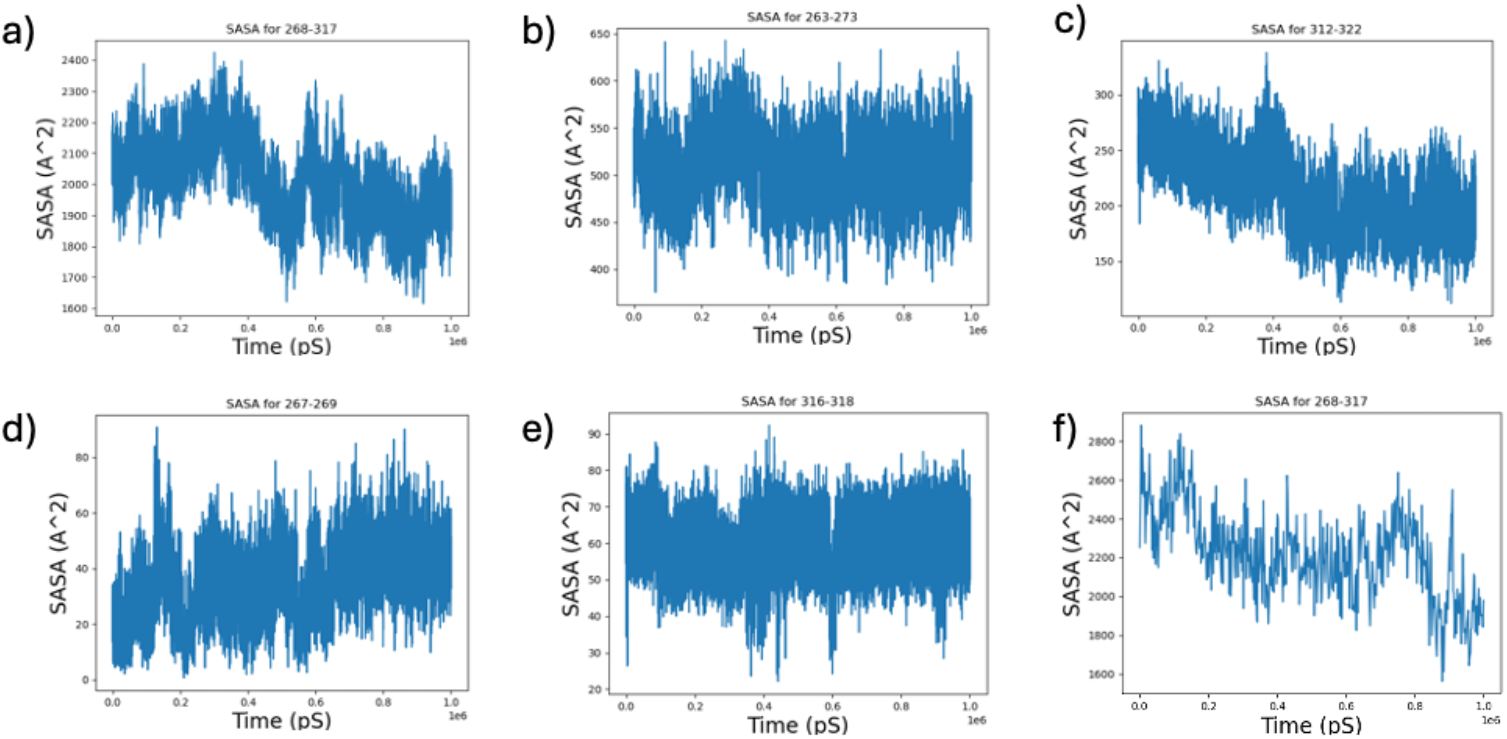
SASA for putative medin cleavage sites along the trajectory of the all atom replicate 1 simulation. a) SASA for the medin region (268-317). b) SASA for the medin N-terminal 10 residue window (residue 263-273). c) SASA for the medin C-terminal 10 residue window (residue 312-322). d) SASA for the medin N-terminal 3 residue window. e) SASA for the medin C-terminal 3 residue window. f) Example of SASA for the medin region (268-317) from a course-grained simulation (500 frame sub-trajectory).

In order to access MfgE8 conformations that may take all atom and coarse-grained simulations too long to sample, an additional set of models of MfgE8 were constructed using CONCOORD *(CONsensus CONfORMational evaluation of COORDinates)*. CONCOORD is a computational tool used for simulating the dynamics of proteins as a computationally cheap pseudo-MD for conformational sampling. It works by employing a combination of MD and distance geometry techniques to explore the conformational space of proteins. CONCOORD was applied to both the AFDB model and the stable compact model generated from the replicate 1 trajectory. During the CONCOORD run, the protein structures explore different conformations outputting an ensemble of 1000 structures for each starting model. The generated ensemble of structures were then used to calculate the SASA values at the putative medin cleavage sites of MfgE8. The SASA values obtained from the CONCOORD models produced a range consistent with the all-atom and coarse-grained molecular dynamics simulations. For the N-terminal compact structure derived CONCOORD models, the SASA obtained values spanned 495-838 (for residues 263-273), while the C-terminal cleavage site (312-322) ranged 220-458. Similarly, the AFDB derived CONCOORD models ranged from 460-896, and the C-terminal SASA values spanned 215-463.

All analyses so far have considered MfgE8 as a globular protein. However, we know that MfgE8 can associate with membranes with previous NMR and mutagenesis studies identifying membrane interacting residues located within 3 spike regions: spike 1 (Tyr23-Trp33), spike 2 (Gln43-Asn47), and spike 3 (Ala78-Gln85) (Ye et al., 2013). Recently Cheng at al (2023) used 150 ns MD simulations enhanced with the highly mobile mimetic model to determine the multimodal binding of Bovine MfgE8 with membranes. The binding occurred primarily via two dynamic structural ensembles: an inserted state similar to previous observations and a previously unreported, highly conserved side-lying state driven by a cationic patch. SASA of the medin cleavage regions in these conformations where MfgE8 was interacting with a membrane were calculated utilising snapshots of both states from simulations conducted by Cheng et al. Again, the SASA results for both were consistent with the all atom MfgE8 in solution simulations, coarse-grained simulations and CONCOORD experiments. For the N-terminal Bovine C2 ‘spike in’ configuration, the SASA was measured at 506, while the C-terminal was 364. Similarly, in the Bovine C2 ‘side on’ configuration, the N-terminal SASA was 600, and the C-terminal was 284.

## Discussion

Despite being the most prevalent localised human amyloid and increasing evidence supporting a role in vascular dysfunction, how medin is cleaved from its parent protein MfgE8 remains unknown. MfgE8 is a promiscuous protein associated with a range of functions and processes (Kamińska et al., 2018). MfgE8 contains three distinct domains with an EGF-like domain forming the N-terminal section, followed by two coagulation domains, with the medin sequence located within the C2 domain. MfgE8 C2 domain is composed of a beta-barrel structure consistent with other coagulation domains (Lin et al., 2007). The molecular dynamics simulations presented here reveal that MfgE8 may adopt a compact conformation with the RGD motif, important for cell attachment and the medin region close in proximity. Given the importance of the RGD motif in the role of MfgE8 mediating uptake of apoptotic cells and the potential pathogenic implications of medin generation we hypothesised that these compact conformations could impact this currently unknown cleavage mechanism. SASA analysis suggests that medin cleavage sites have low accessibility indicating protection from cleavage. Additionally, formation of compact structures observed in the simulations result in a further decrease in accessibility of the medin region and C-terminal 10-residue window, consistent with these conformations protecting against cleavage mechanisms. Interestingly, analysis of a 3-residue window surrounding the putative medin cleavage sites did not show a reduction in accessibility upon formation of the compact structure. This suggests that a greater area of either a 10-residue window or the entire medin region may be needed to influence accessibility and therefore potential cleavage resulting in medin generation from MfgE8. As expected, modelling of MfgE8 in the presence of an integrin dimer shows the RGD motif in contact with integrin (data not shown) preventing formation of the previously observed compact MfgE8 formation.

The sequence of medin has been described as ragged with minor species also identified to have start sites either side of the consensus 50 amino acid sequence (Häggqvist et al., 1999). These include residues at positions 264 and 273 of MfgE8. Our 10-residue window analysis for SASA incorporated these start sites covering residues 263-273 with no obvious accessibility favouring cleavage at either of these alternative start sites in addition to arginine at position 268.

Given the apparent protection from cleavage rather than resulting increase in accessibility over time as expected we hypothesise that something must change that initiates the accessibility to these sites necessary for medin production. One obvious condition that the current study has not incorporated is the presence of post-translational modifications. Specifically, this study does not account for the glycosylation state of known sites on MfgE8. MfgE8 is known to be glycosylated at a range of positions including asparagine residues at positions 228, 238, 325, 329 and 350 within the C2 domain (Picariello et al., 2008). Annotating these modifications onto the MfgE8 domain structure reveals a cluster of N-glycosylation sites around the sequence of medin (Figure 1a). Glycosylation is critical for protein folding and various other cellular processes (Ruotsalainen et al., 2022). Additionally, N-glycosylation has recently been described to be protective against protein aggregation due to steric hindrance, with overrepresentation reported at the N terminus of aggregation prone regions (APRs) (Duran-Romaña et al., 2024). Considering glycosylation in the modelling of MfgE8 could provide a more comprehensive understanding of both inter- and intra-domain interactions, as well as accessibility to the putative medin cleavage sites. We note that known glycosylation sites of MfgE8 are only present in the medin C2 domain, despite the similar sequence identity and structure with the C1 domain -sequence similarity of 97.5%.

The medin region lies between the Asn238 and Asn325 glycosylation sites. In experimental structures and AFDB models, Asn238 is located within a loop that is in close proximity to the N-terminal medin cleavage site (as can be observed in Figure 12). A recent Finnish study (Ruotsalainen et al., 2022) has uncovered a protective association against atherosclerosis with certain MfgE8 variants. Notably, the sequence mutations from one variant (rs534125149) are localised to the aforementioned loop region within the C2 domain, in proximity to both the N-terminal medin cleavage site and approximately 20 Å away from key amino acids involved in membrane binding. The insertion of an additional asparagine residue within this loop shifts the sequence of the Asn238 glycosylation site from NNS to NNNS in this variant. This could result in impaired or disrupted glycosylation, or it could also shift the site from Asn238 to Asn239, given the consensus sequence for glycosylation is described as NxS (Rao & Wollenweber, 2010). The insertion variant could also affect the interaction of this loop region with the N-terminal start site of medin (Figure 13). In addition, this loop region is interacting with the cell membrane in the side lying conformation of the C2 domain previously reported (Cheng et al., 2023). Therefore, the insertion and its potential impact on glycosylation might affect the ability of the C2 domain to adopt a side lying conformation.

**Figure 13:**
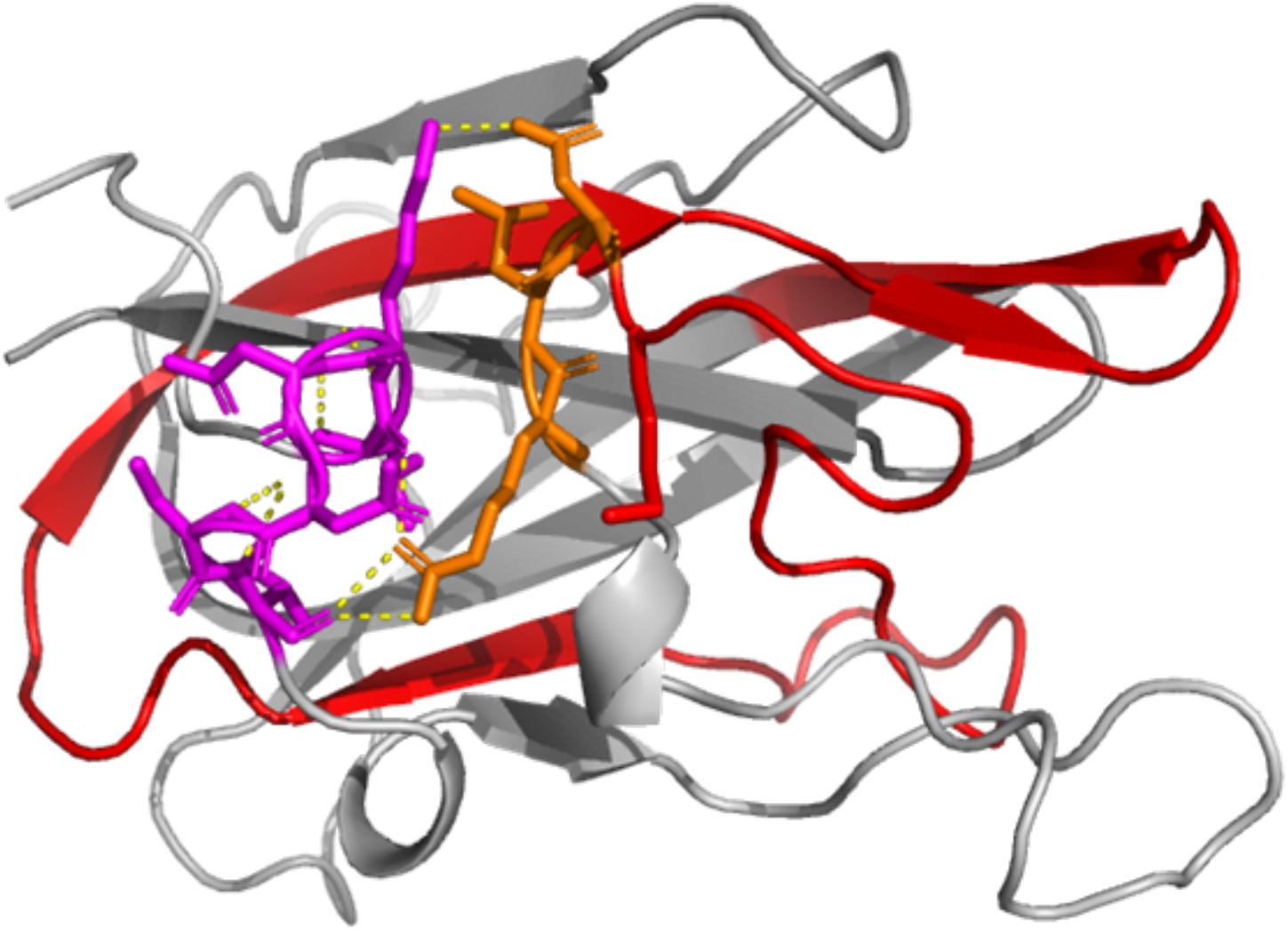
C2 domain of MfgE8 with medin region highlighted in red. Side-chains of N-terminal start-site of medin (residues RLD) shown in orange, loop containing the N238 (LKNNSI) shown in purple. Hydrogen bonds computed with pymol shown in yellow.

Medin accumulation within blood vessels has been shown to be age-dependent, occurring in nearly all Caucasians studied over the age of 50 (Mucchiano et al., 1992). Alterations in glycosylation are also linked to the aging process and associated with response to immune stimuli and other processes that can become dysregulated upon aging (Cindrić et al., 2021). Consequently, alterations in glycosylation patterns have been proposed as potential serum/plasma biomarkers in a range of age-related diseases (Cindrić et al., 2021; Reily et al., 2019).

Exploration of modelling and dynamics taking into account the presence of this post-translational modification and others as well as sequence variants might shed light on the mechanisms by which the medin cleavage sites become accessible to proteolysis and any potential age-related association with this process. The recent release of the cutting-edge modelling tool AlphaFold3 (Abramson et al., 2024), combines a residue-based representation of amino acids with an atomic representation of all other groups to model assemblies containing proteins with covalent modifications given their sequences and chemical structures. Utilisation of this software combined with MD simulations will provide a state-of-the-art platform to understand how glycosylation of key residues impacts the conformation of MfgE8 as well as the accessibility of the proposed medin cleavage sites.

## Conclusions

MfgE8 exhibits a propensity for adopting a compact conformation facilitated by interdomain electrostatic interactions, thereby promoting stability. Interestingly, adopting this compact conformation appears to decrease the solvent-accessible surface area of the medin region and the C-terminal cleavage site. To enhance accessibility to the medin cleavage sites of MfgE8, alterations in its local environmental conditions may be necessary where the protein unfolds and becomes more prone to cleavage.

## Funding Acknowledgment

US National Institutes of Health/National Institute of Aging R21AG075543.

